# Sleep deprivation and the rodent psychomotor vigilance test (rPVT): Assessing lapses in attention, the response stimulus interval effect, and time on task in rats using food reinforcement

**DOI:** 10.1101/2024.04.10.588729

**Authors:** Catherine M. Davis, Victoria Elliott, Joan Smith

## Abstract

**Study objective:** Sleep deprivation (SD) impairs sustained attention when assessed with the psychomotor vigilance test (PVT). Food restriction attenuates the effects of SD on sustained attention in rats, possibly limiting translation of rodent tests. Our goal was to determine if an rPVT requiring high baseline performance was sensitive to the effects of SD when using food restriction and reinforcement.

**Methods:** rPVT-trained rats experienced SD and were tested the following day.

**Results:** SD significantly increased lapses, and this effect was specific to shorter response-stimulus intervals. Decreased percent correct responses and increased slow reaction times were also found following SD.

**Conclusion:** The rPVT is sensitive to the performance impairing effects of SD in food restricted rats, a common methodology used to train and maintain performance on operant tests.

## Introduction

Sleep loss reduces sustained attention, degrades various cognitive domains, and leads to functional impairment (Hudson et al., 2020). The human PVT is the “gold standard” objective measure of sleep loss-induced changes in sustained attention, psychomotor speed, impulsivity, and state stability (Basner & Dinges, 2011). In this test, subjects must monitor the location of a randomly occurring stimulus; when it appears, they must make a response as quickly as possible. Several rodent versions of the PVT (rPVT) have been developed (Christie et al., 2008; Davis et al., 2015; Davis et al., 2014; Deurveilher et al., 2015; Oonk et al., 2015). We previously compared performances on each version, and noted that high performance levels (e.g., high percentage of correct responses, low numbers of premature responses) provided good baselines to assess drug and stressor effects (Davis et al., 2016). Importantly, food restriction and reinforcement appear to attenuate the effects of SD on attention tests in rodents (Loomis et al., 2020), with many studies utilizing water restriction and reinforcement for this reason.

We developed a version of the rPVT using food restriction and reinforcement that requires rats to maintain high levels of baseline performance (Davis et al., 2015; Davis et al., 2014; Davis et al., 2016), and previously reported robust response-stimulus interval (RSI) and time-on-task effects where healthy, non-sleep-deprived rats displayed predictable changes in performance (Davis et al., 2016). The goal of the current study was to determine if this version of the rPVT is sensitive to the performance-impairing effects of SD.

## Methods

### Subjects and Apparatus

Animal care was conducted according to Public Health Service (PHS) Policy and the Institutional Animal Care and Use Committee of the Johns Hopkins University and Uniformed University of the Health Sciences approved all procedures. Both programs are accredited by the Association for the Assessment and Accreditation of Laboratory Animal Care (AAALAC). Male Long-Evans rats (N=4, Envigo, East Millstone, NJ) were received at approximately 10–12 weeks of age, singly housed in plastic cages with enrichment toys, maintained on a 12:12 h light-dark schedule (lights on at 0600, Zeitgeber time [ZT] ZT0), and at an ambient temperature of 23°C. Rats were maintained at 90% of their free-feeding weights by being fed measured amounts of chow each day after they performed the rPVT. Water was freely available in the home cage. Rats were run in 30-min rPVT sessions at the same time each day (1100; ZT5). Operant chambers contained one nose-poke key, cue and house lights, and a food cup for delivery of pellets, and were enclosed in sound-attenuating cubicles equipped with exhaust fans. MedPC® IV programs controlled experimental contingencies; all data was recorded on a trial-by-trial basis.

### Rodent Psychomotor Vigilance Test (rPVT)

Rats were trained to respond on a nose-poke key for food. Once acquired, rPVT training began. Sessions began with the onset of the house light. After a variable delay of 3–10 sec, the nose-poke key was illuminated. A correct response was defined as a response on the nose poke key within 2 sec after the light onset (i.e., 2-sec limited hold, LH) and was reinforced with a pellet. Responses prior to the light onset (premature response) were not reinforced and punished with an 8 sec time out, while responses after the 2-sec interval had elapsed or no response (omission) were not reinforced. The delay period for the next trial began after a 1-sec inter-trial interval, timed either after the response or the end of the 2-sec LH, whichever occurred first. Data collected were the number of correct responses (“hits”), premature responses, omissions, and lapses in responding (omissions plus correct responses greater than twice each rat’s mean response latency for that session). Summary measures were expressed as total numbers and percentages. Response latencies were recorded in milliseconds, and summarized by calculating various reaction times (e.g., 10^th^ percentile, median, 90^th^ percentile, mean). Each performance measure was also calculated based on the response stimulus interval during which the measure occurred or based on time-on-task (i.e., performances binned into 6-minute intervals across the 30-min session time). Criteria for inclusion in this study were >75% percent correct and < 25% premature responses for four out of the five daily test sessions during one to two weeks prior to any manipulation.

### Acute sleep deprivation

Rats were moved from their home cages to sleep fragmentation chambers (Model #80391, LaFayette Instrument, LaFayette, Indiana), which are standard home cages containing an automated sweep bar that moves across the chamber. Sweep duration can be set at different intervals, and was set to ‘continuous’, where the bar moves across the chamber once every 7.5 seconds. Rats were housed in these chambers for two days with the sweep bar stationary as baseline days to ensure that housing alone in this chamber did not impair rPVT performance. Sweep bars were turned on starting at lights off (1800; ZT12) and were run continuously throughout the lights off period and the beginning of the lights on period of the next day until each subject was moved to their rPVT chamber for testing (ZT5). Rats were then returned to standard home cage housing. Two bouts of SD were completed, separated by approximately one month, and the two bouts were averaged together for data analysis.

### Data analysis

Paired t-tests were used to assess the effects of SD on the number of lapses, percent correct responses, median reaction time, mean reaction time, mean speed, Q90 reaction times, hits, long RT hits, and omissions. To assess the response-stimulus interval and time-on-task effects, separate two-way repeated-measure ANOVAs were used, followed by Bonferroni’s multiple comparisons test. Statistical analyses were completed using GraphPad Prism (v10.2.0). Alpha was set to 0.05 for significant effects.

## Results

Acute SD resulted in a significant increase in the number of lapses [t(3)=10.05, *p* = 0.0021, *r*^*2*^ = 0.9711; Figure 1A] and significantly greater percentage of lapses were emitted at shorter response-stimulus intervals [RSI; response stimulus interval x sleep day interaction: F(6,18) = 10.04, *p* < 0.0001], with significant increases in lapses emitted at intervals from 3-8 seconds in duration (all p’s < 0.0476; Figure 1C). The greatest increase in lapses were at the two shortest RSIs (3-4 sec and 4.1-5 sec). Significant main-effects of time-on-task and SD were found on the percentage of lapses emitted [F(4,12)=24.19, *p* < 0.0001 and F(1,3)=36.29, *p* = 0.0092; Figure 1D], but these did not vary by time bin; thus, SD increased lapses equally across the PVT session. SD significantly decreased percent correct responses [t(3)=14.70, *p* = 0.0007, *r*^*2*^ = 0.9863; Figure 2A], and significantly increased Q90 reaction times (the slowest 10^th^ percentile) [t(3)=4.502, *p* = 0.0205, *r*^*2*^ = 0.8711; Figure 2B].

**Figure 1.**
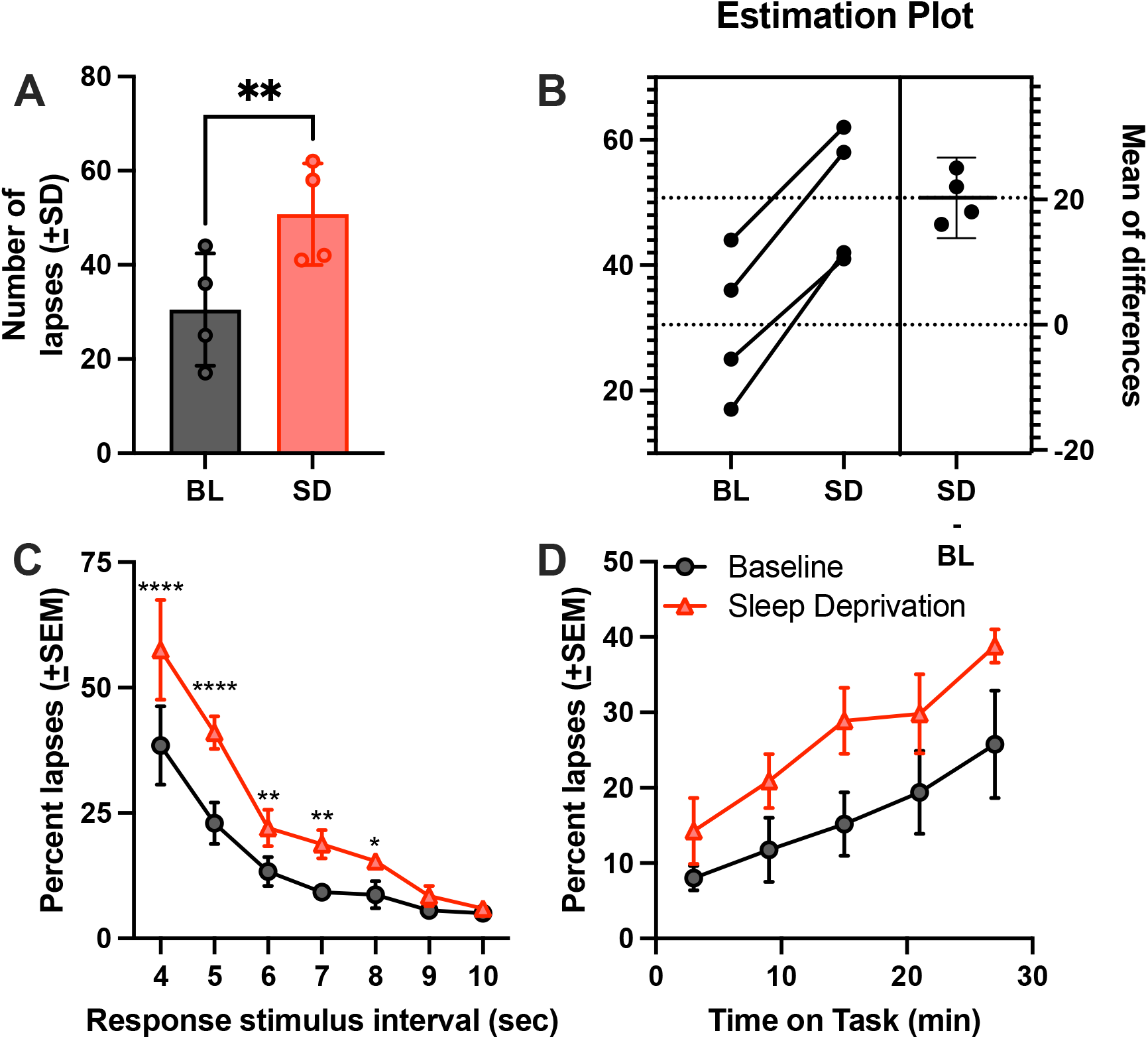
Sleep deprivation increases lapses on the rPVT when responding is reinforced with food pellets. A) Mean number of lapses increased following acute sleep disruption (SD) compared to baseline (BL). B) Estimation plot showing mean difference (effect size). Error bars indicate 95% CI of the mean difference. C) Sleep deprivation significantly increased lapses at response stimulus intervals between 3-8 seconds. RSI 4 = RSIs from 3 – 4 sec, RSI 5 = RSIs from 4.1 – 5 sec, etc. D) Lapses increase across time on task, but not at specific time bins following sleep deprivation \**p*<0.05, \*\**p*<0.01, \*\*\*\**p*<0.0001.

**Figure 2.**
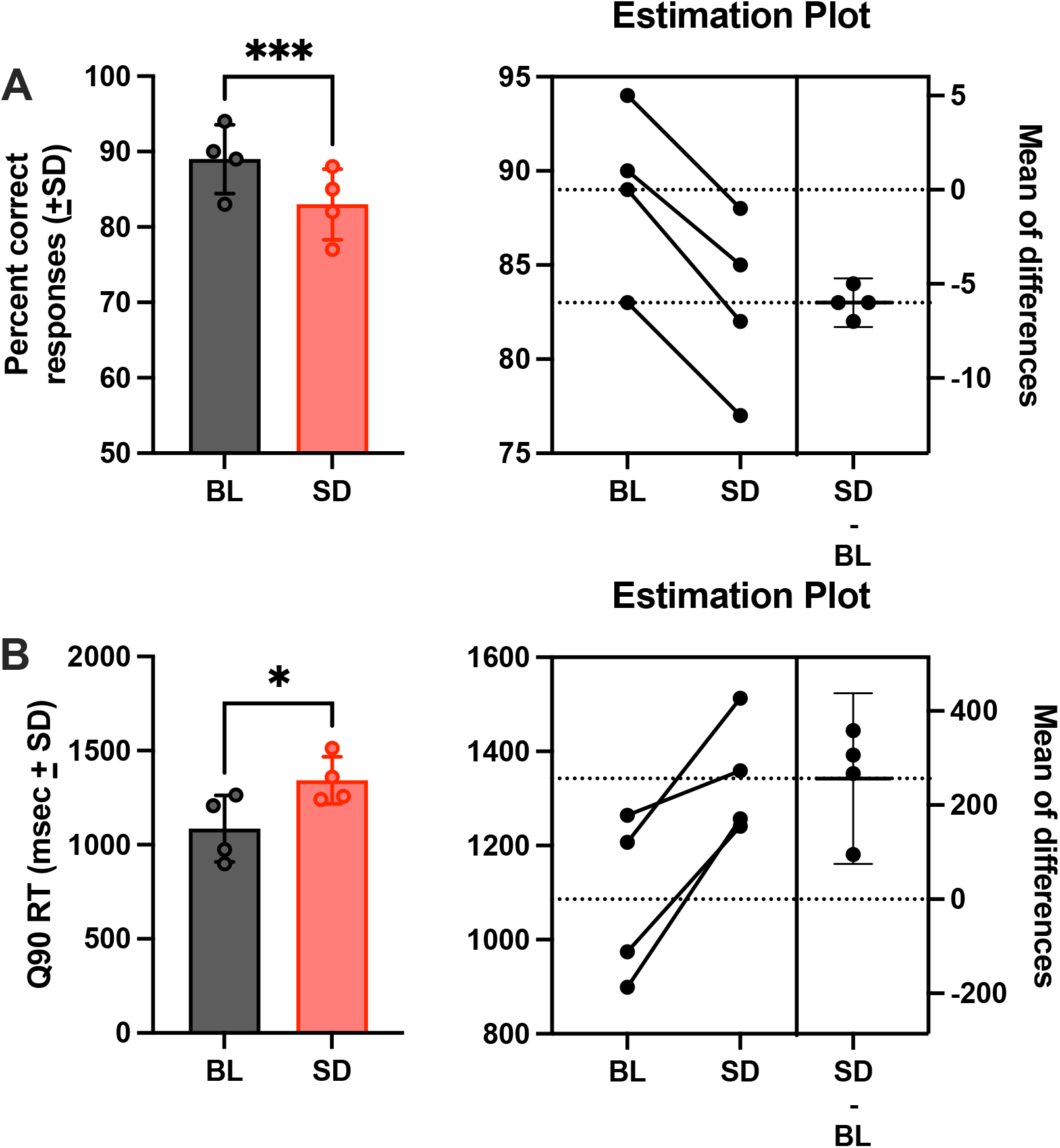
Sleep deprivation decreases percent correct responding and increases the slowest 10^th^ percentile reaction times (Q90). A) Percent correct responses were significantly decreased following sleep deprivation (SD) when compared to baseline (BL), however, rats still maintained > 75% correct responding following sleep deprivation. B) Sleep deprivation significantly increased Q90 reaction times, with an average increase of ∼300 msec. Estimation plots are included to show mean difference (effect size) and 95% CIs. \**p*<0.05, \*\*\**p*<0.001.

SD decreased percent correct responses at specific RSIs [RSI x SD interaction: F(6,18)=6.267, *p* = 0.0011], with fewer correct trials found at the shortest RSIs (3-4 and 4.1-5; all *p*’s < 0.0035; Figure 3A). However, even after SD, animals continued to emit a high number of correct responses (“hits”), even though this was significantly decreased from baseline [t(3)=4.392, *p* = 0.0219, *r*^*2*^ = 0.8654; Figure 4A]. SD significantly increased omissions [t(3)=5.081, *p* = 0.0147, *r*^*2*^ = 0.8959; Figure 4C] and trended to increase correct trials with long response times [t(3)=2.717, *p* = 0.0727; Figure 4B]. Thus, animals continued to perform at high levels even after SD, which suggests rats did not fall asleep during the rPVT session. SD did not affect mean or median reaction times, mean speed, Q10 reactions times, or premature responses (all *p*’s > 0.2594).

**Figure 3.**
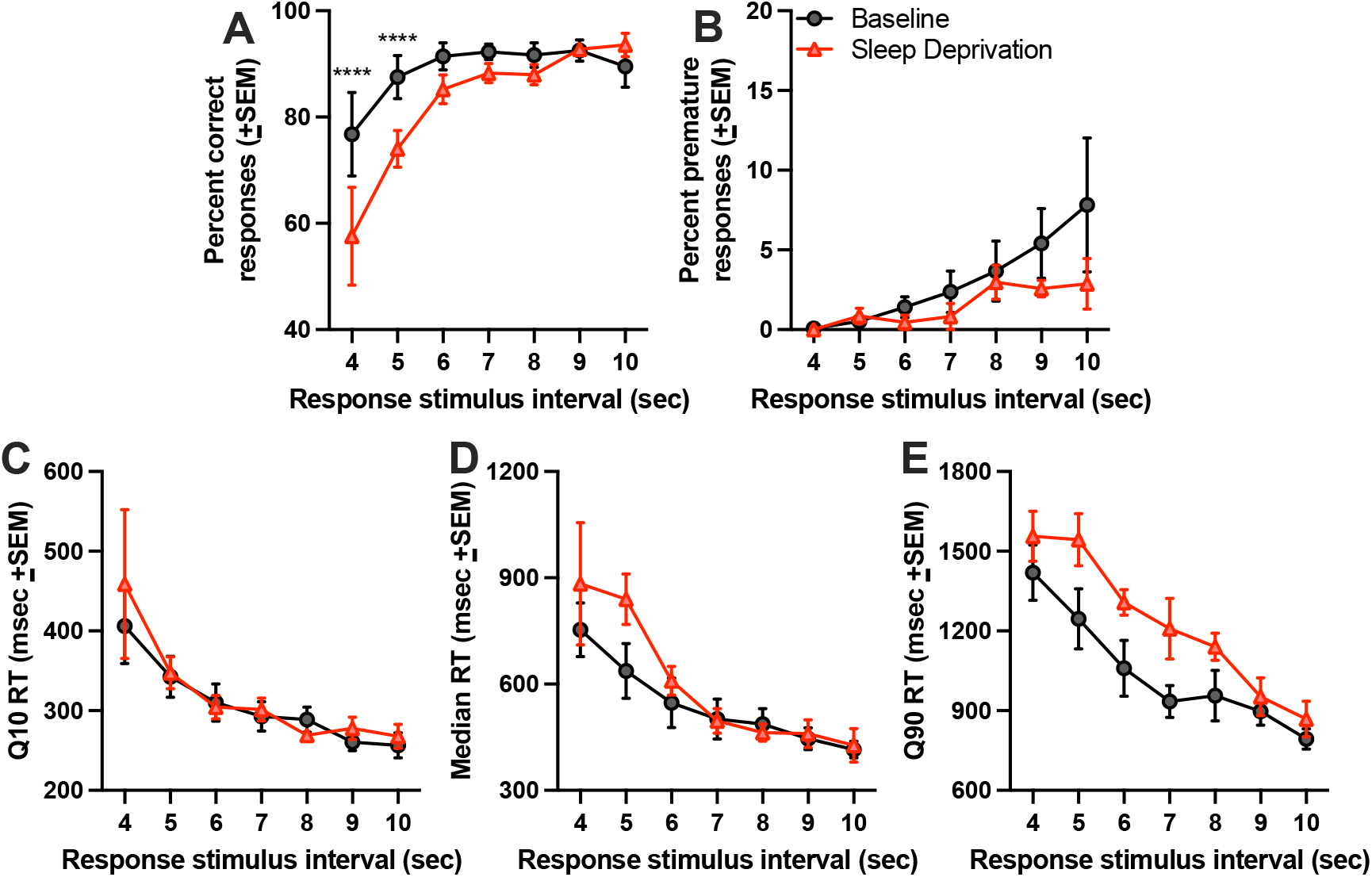
Sleep deprivation decreases percent correct responses at the shortest RSIs. A) Mean percent correct responses at each RSI on the rPVT at baseline and following sleep deprivation. SD significantly decreased percent correct responses at the two shortest RSIs. B) Mean percent premature responses. C) Mean Q10 reaction times. D) Median reaction times. E) Mean Q90 reaction times significantly differed from baseline overall (see Fig 2), but not in an RSI-dependent manner. \*\*\*\**p*<0.0001.

**Figure 4.**
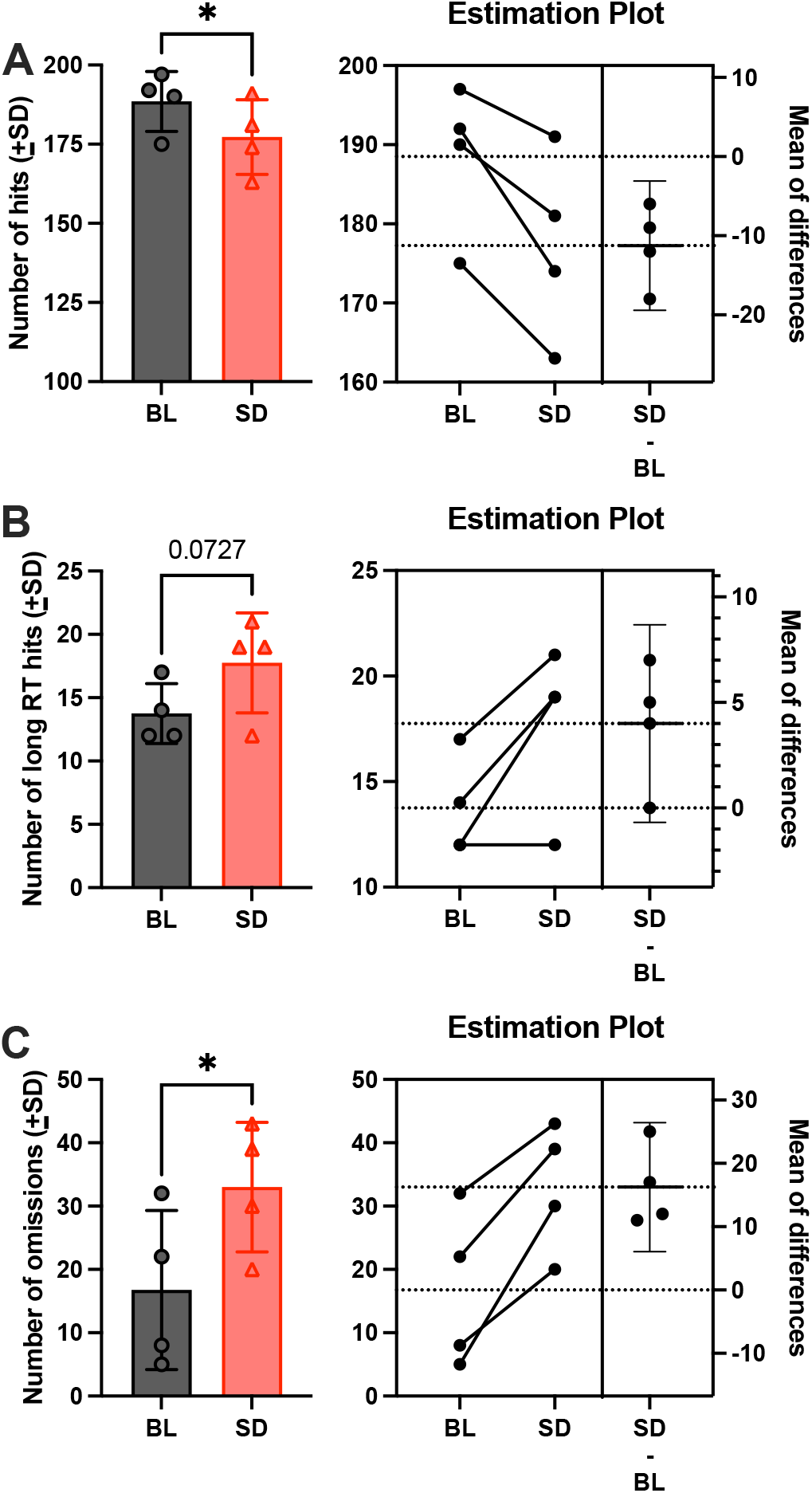
Sleep deprivation increases slow response trials and misses on the rPVT, which leads to decreased number of hits. A) Mean total hits (correct trials) on the rPVT significantly decreased following sleep deprivation (SD) compared to baseline (BL), even though the total number of hits was still high. B) Sleep deprivation trended to increase ‘long hit trials’, which are trials where animals took twice their mean reaction time to respond correctly (one trial type included in lapses). C) Sleep deprivation significantly increased the number of missed trials where no response was recorded (omissions; the second trial type included in lapses). Total responses in B and C are added together to calculate total number of lapses (see Fig. 1). Estimation plots are included to show mean differences (effect size) and 95% CIs. \**p*<0.05.

## Discussion

We found that SD impaired sustained attention in food-restricted rats performing a food-reinforced rPVT. Specifically, SD significantly increased the number of lapses emitted, and these effects were more robust at shorter RSIs, with equivalent numbers of lapses emitted at long RSIs (9-10 sec) between baseline and SD. The most lapses were emitted at the two shortest RSIs, which was accompanied by a significant decrease in percent correct responses, suggesting that lapses following SD during these short RSIs consisted primarily of omissions. SD also slowed the slowest reaction times (Q90 RTs). Thus, following SD, rats had more difficulty responding to quickly occurring stimuli. Further, these results support the translational nature of the rPVT, with the number of lapses increasing following SD in a manner similar to humans (Basner & Dinges, 2011). Taken together, these data demonstrate that the food-reinforced rPVT is sensitive to the performance-impairing effects of SD.

Loomis and colleagues (2020) recently reported that food restriction results in functional resilience to SD when compared to *ad lib* feeding conditions using a simple response latency task (SRLT), where rats are required to monitor the location of a randomly occurring light stimulus. The rPVT and SRLT are similar, but there are important methodological considerations that likely increased sensitivity of the rPVT to the impairing effects of sleep deprivation in food-restricted rats. First, baseline performance on the rPVT was high, with rats displaying an average of 89% correct responding (Figure 2A). In contrast, food-restricted rats performing the SRLT emitted significantly greater premature responses than subjects under *ad libitum* conditions, even though both groups emitted similar numbers of correct trials and omissions. Thus, regardless of the test, rats with a higher number of correct trials and fewer premature responses are more sensitive to the effects of SD. These data suggest that baseline performance is an important factor for whether the effects of SD are detectable in rodent sustained attention tests. While the rPVT measures lapses, which are a combination of correct trials with slow response times and omissions, the SRLT measures omissions only. Despite these differences, SD significantly increased omissions on the SRLT under *ad libitum* food conditions and the rPVT using food restriction (Figure 4). Significant time-on-task effects were found for lapses, percent correct responses, and Q90 reaction times for the rPVT, and SD increased these measures overall (but not in a time bin-dependent manner). These effects are similar to those reported by Loomis and colleagues (2020), in addition to previous rPVT reports employing water restriction and reinforcement (Christie et al., 2008; Deurveilher et al., 2015; Oonk et al., 2015). Finally, while SD did increase Q90 reaction times on the rPVT, it did not significantly affect mean, median, or Q10 reaction times. These results are similar to those found in food-restricted rats performing the SRLT, where SD did not change median response times.

There are several limitations to the current study. One is the lack of electroencephalogram and electromyogram (EEG/EMG) recordings, which would have shown the effects of SD on physiological measures of sleep, in addition to how these processes might be affected by food restriction. A small group of subjects was used in this study, and while many of the baseline effects (e.g., RSI, time-on-task) were the same as those we reported previously in a large cohort of rats (Davis et al., 2016), a larger group size will be needed to replicate and extend these findings. Future studies should also examine both sexes. Finally, the lack of an effect of SD on mean and median reaction time measures, and mean speed, could result from food restriction, but more work is needed in a larger group of rats to assess this possibility.

In conclusion, the rPVT using food reinforcement in food restricted rats is sensitive to the effects of a single night of SD, with significantly increased lapses, increased Q90 reaction times, and decreased percent correct responses. These data demonstrate the usefulness of the rPVT for investigating the attention-impairing effects of SD in rats.

## Acknowledgements

We would like to thank Dr. Robert Hienz for discussions related to design and completion of this study, and Ray Smith, Blanca Bravo, and Stacey Perry for their technical assistance. This work was supported by Space@Hopkins seed grant funding (to CMD) and NASA NNX15AC71G and 80NSSC22K0022 (to CMD). Some of the authors are employees of the U.S. Government, and this work was prepared as part of their official duties. Title 17 U.S.C. §105 provides that ‘Copyright protection under this title is not available for any work of the United States Government.’ Title 17 U.S.C §101 defines a U.S. Government work as a work prepared by a military service member or employees of the U.S. Government as part of that person’s official duties. The opinions and assertions expressed herein are those of the author(s) and do not reflect the official policy or position of the Uniformed Services University of the Health Sciences, the Armed Forces Radiobiology Research Institute, Department of the Navy, the Department of Defense, or the US Federal Government.

