## Supplementary information for "Sleep deprivation and the rodent psychomotor vigilance test (rPVT): Assessing lapses in attention, the response stimulus interval effect, and time on task in rats using food reinforcement"

**Methods:** rPVT percentages were calculated as follows: correct responding = number of correct responses / (corrects + premature responses + omissions); premature responding = number of premature responses / (corrects + premature responses + omissions); lapses = (omissions + long RT hits) / (corrects + premature responses + omissions).

**Results:** Significant main-effects of time-on-task effect and sleep deprivation were found for percent correct responses [$F(4,12)=30.66$, $p < 0.0001$ and $F(1,3)=133.6$, $p = 0.0014$; Figure S1A], and Q90 reaction times [$F(4,12)=5.075$, $p = 0.0125$ and $F(1,3)=13.07$, $p = 0.0364$; Figure S1B]; no time-on-task x sleep deprivation interactions were found ($p \geq 0.1464$), which supports the finding that sleep deprivation decreased percent correct responses and increased Q90 RTs equally across the rPVT session.

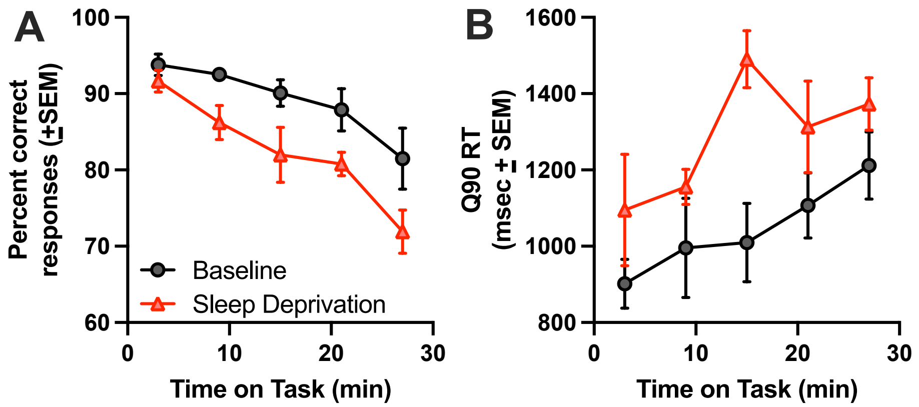


**Figure S1. Time on task effects for percent correct responses and Q90 reaction times (RT) for the rPVT.** A) Percent correct responses decrease across time on task following sleep deprivation, but not differently at specific time bins. B) Q90 reaction times increase across time on task following sleep deprivation, but not differently at specific time bins.
